# Laying the Foundation: Modern Transformers for Gold-Standard Sleep Analysis and Beyond

**DOI:** 10.1101/2024.01.18.576246

**Authors:** William G. Coon, Mattson Ogg

## Abstract

Accurate sleep assessment is critical to the practice of sleep medicine and sleep research. The recent availability of large quantities of publicly available sleep data, alongside recent breakthroughs in AI like transformer architectures, present novel opportunities for data-driven discovery efforts. Transformers are flexible neural networks that not only excel at classification tasks, but also can enable data-driven discovery through un-or self-supervised learning, which requires no human annotations to the input data. While transformers have been extensively used in supervised learning scenarios for sleep stage classification, they have not been fully explored or optimized in forms designed from the ground up for use in un-or self-supervised learning tasks in sleep. A necessary first step will be to study these models on a canonical benchmark supervised learning task (5-class sleep stage classification). Hence, to lay the groundwork for future data-driven discovery efforts, we evaluated optimizations of a transformer-based architecture that has already demonstrated substantial success in self-supervised learning in another domain (audio speech recognition), and trained it to perform the canonical 5-class sleep stage classification task, to establish foundational baselines in the sleep domain. We found that small transformer models designed from the start for (later) self-supervised learning can match other state-of-the-art automated sleep scoring techniques, while also providing the basis for future data-driven discovery efforts using large sleep data sets.

## I. Introduction

Sleep is fundamental for good health and optimal performance, and its accurate assessment is critical to sleep medicine and research. Sleep assessment (“sleep scoring”) has traditionally been done by human experts trained to classify 30-second segments (epochs) of sleep into one of five stages according to patterns discernible principally in the electroencephalogram (EEG)^†^ [2, 3]. Modern neural network architectures, supported by the recent availability of large quantities of publicly available sleep data, are beginning to supplant this tedious, manual approach, having met and exceeded human expert performance [1, 4, 5]. Successful approaches span the gamut of developments in the previous 15 years of AI research, including convolutional and recurrent neural networks[4–6], U-nets [7], graph networks [8], resnets [9], and, increasingly, a wide variety of attention- and transformer-based architectures [10–15]. Today, the question is not whether automated approaches can serve instead of manual scoring, but rather how far they can be pushed to discover latent, heretofore unutilized health information from sleep data in a principled, data-driven way (two excellent recent examples are a study by Decat et al. [16], and a recent effort to update traditional sleep staging with a “high-frequency” classifier [7]). Generating foundation models through self-supervised learning is a promising a way forward in this respect.

One definition of a foundation model is a neural network architecture that can be pre-trained, often using unlabeled data and self-supervised training, whose embeddings of the data space are so rich as to allow for multiple future use cases, often via efficient fine-tuning on a specific application afterward. Pre-training vastly accelerates learning specific tasks, providing the practical advantage that multiple model applications can be derived from the same pretrained model. Furthermore, since they encode compact representations of the data at such high fidelity, foundation models are particularly well-suited for data-driven discovery. They can be flexibly applied to many different ends (ex. classifying sleep stages, apnea state, disease diagnosis, age, etc.), or be interrogated to discover new information embedded within (for example, via naïve clustering of its embeddings, which can then be correlated with health or examined for implicit structure). These methods offer new ways to quantify and correlate sleep signals data with the many health states and conditions known to interact with sleep.

In pursuit of these goals, model selection is crucial since performance comparisons will require appropriate baselines, and these in turn require using the same architecture in supervised and self-supervises cases. While there is an abundance of sleep stage classifiers that leverage self-attention and transformers (see [10–15] for recent examples), it is not clear which of these would be best for self-supervised learning followed by fine-tuning; none have been designed from the bottom up with self-supervised learning in mind. One way around this is to begin with a model already known to perform well in self-supervised applications, and determine which optimizations are needed for that architecture to perform the relevant baseline task, i.e.., supervised sleep stage classification. Accordingly, we explored optimizations to a transformer-based neural network model developed for acoustic speech processing (“HuBERT”) [17] and language modeling [18] which have demonstrated track records of successful self-supervised learning. Speech processing and language modeling are both sequence learning tasks with similarities to mapping sleep EEG signals to a sequence of sleep stage labels, so we reasoned that these models should be able to learn the canonical 5-class sleep stage classification task. A detailed analysis of how these models can be used for traditional sleep-staging provides a valuable baseline for future work in foundation model building. To this end we examined different configurations of model size, input data size (sleep sequence length), input channels, and the use of multiple or single nights of sleep during training. Here we report our findings and discuss transitioning the trained models to analyze sleep on wearable devices (useful for generating data at-scale) and to enable data-driven discovery using foundation model variants in future work. One such example using self-supervised learning is discussed in this work’s companion paper, appearing in this issue of EMBC Proceedings as Ogg & Coon 2024.

### II. Methods

### A. Data and Preprocessing

We compiled a large corpus of training data from the National Sleep Research Resource [19]. We obtained data from the Multi-Ethnic Study of Atherosclerosis (MESA [20]), the MrOS Sleep Study[21], the Sleep Heart Health Study [22], the Wisconsin Sleep Cohort [23], the Nation-wide Children’s Hospital Sleep DataBank (NCHSDB) [24], the Study of Osteopathic Fractures (SOF) [25], and the Cleveland Family Study (CFS) [26]. This comprised data from over 11,000 nights of sleep from over 9,500 individuals. We held out data from 915 participants (or approximately 10%) to monitor training. Two additional datasets were obtained for external validation and testing (i.e., after training and using held out participants for internal validation, we evaluate final perfomance on separate hold-out collections whose data the model had not encountered during training). We used the Dreem Open Datasets (both the Healthy and OSA cohorts) [27] for external validation and the Sleep-EDF Database Expanded dataset [28] as a test set.

Each sleep session (i.e., each EDF^‡^ file) underwent the following pre-processing regime. We retained one central EEG recording (i.e., C3 or C4 from the International 10-20 System) and one (left or right) EOG recording (any sleep session without these channels explicitly marked was excluded). These channels were re-sampled to a 100Hz sampling rate and then robustly re-scaled by subtracting the timeseries’ medians and dividing by the difference between each’s 25th and 75th percentiles (thus achieving an inter-quartile range (IQR) of 1 for each channel as in Perslev et al.[7]). Any values exceeding a value of *±*20 median absolute deviations were clipped to remain within that range. No further signal processing to the input data was applied. For the Sleep-EDF dataset, we used the Fpz-Cz and Horizontal EOG channels, and truncated the large awake periods to just the 30 minutes preceding and following sleep.

Each 30-second epoch of EEG and EOG data, with their corresponding sleep labels, was then extracted based on the recording’s annotations. Any epochs that were not explicitly labeled with a sleep stage, or were labeled as artifacts, were removed. The remaining epochs were then grouped into sequences of 21 (10.5 minutes) or 101 (50.5 minutes) consecutive epochs^§^. These sequences of labels along with their corresponding time series data (at a 100 Hz sampling rate) served as the training labels and input data for the transformer model, respectively. We over-sampled the training and internal validation data by a factor of four when sequencing epochs (21-epoch sequences were generated every 5 epochs; 101-epoch sequences were generated every 25 epochs). External validation and test data were not over-sampled (101- and 21-epoch sequences were generated with no overlap).

### B. Network Architecture and Training

Similar to work in the speech and audio domain [17], our model learns to map time series input to a sequence of labels, which in our case correspond to one of 5 sleep stages (W, N1, N2, N3 or REM). Transformers are an excellent choice for this application because of their ability to model long-distance temporal dependencies within the data.

The model architecture begins with a series of 1-dimensional convolution layers as a front end, to learn a set of filters directly from the input time series (as in previous work [4, 5]). A layer-norm was applied between the convolution and activation functions in each layer. This front end was followed by a linear projection layer, positional encoding, a stack of transformer encoder layers, temporal averaging (to align the higher temporal resolution of the convolution/transformer layers with epoch labels every 30-seconds), another linear projection layer, and, finally, an output layer. We use gelu activation functions after every convolution layer and within the transformer blocks. Convolution and linear layer weights were initialized with a Kaiming normal function. This common architecture is depicted in Figure 1, and more details on our variants of these models can be found in Figure 2.

**Figure 1.**
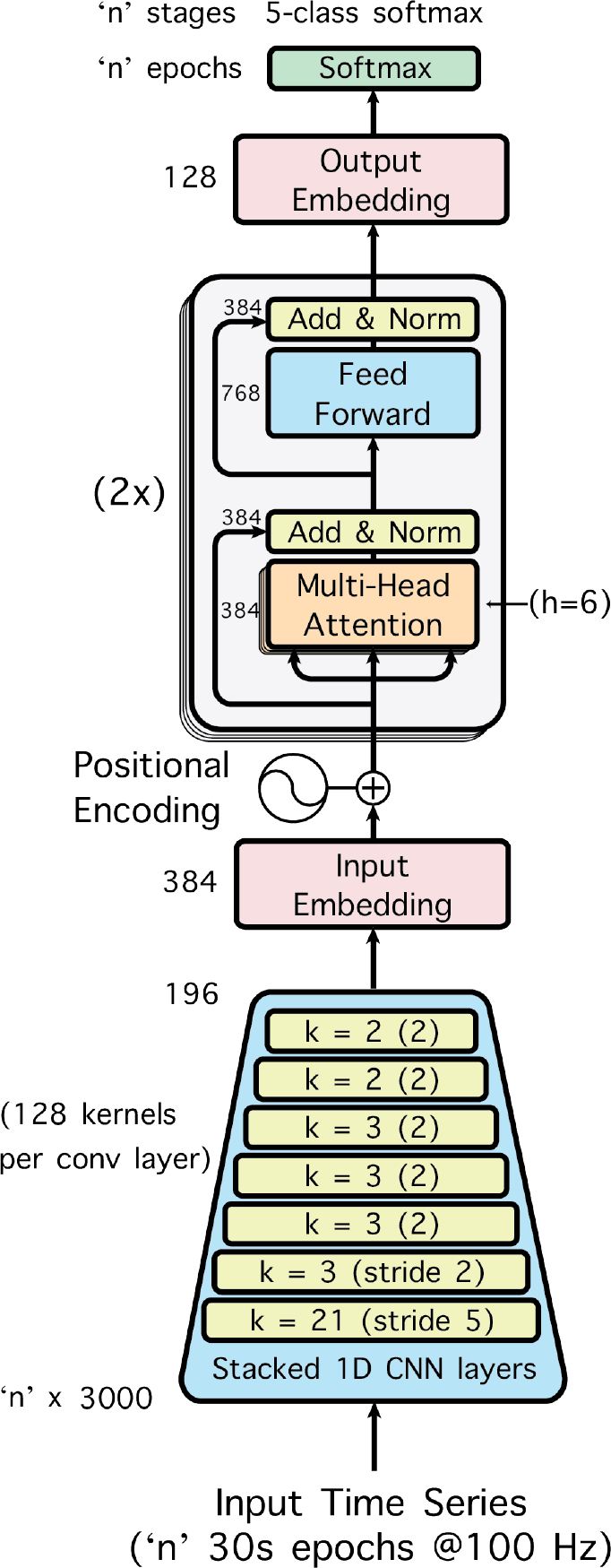
Transformer-based sleep staging model. The model is trained to map input time series data to a sequence of sleep stages. A stack of 1D CNNs first extracts features from the raw time series, and is followed by a linear projection to an input embedding, positional encoding (as in Vaswani et al., 2017 ([29])), multiple transformer encoder layers, another projection to an output embedding, and a final softmax layer for classifying sleep-stage labels (Wake, N1, N2, N3, REM) for each 30-second epoch in the input time series. The parameters needed to exactly reproduce this in PyTorch are included in the figure, here showing the configuration of the small model variant.

**Figure 2.**
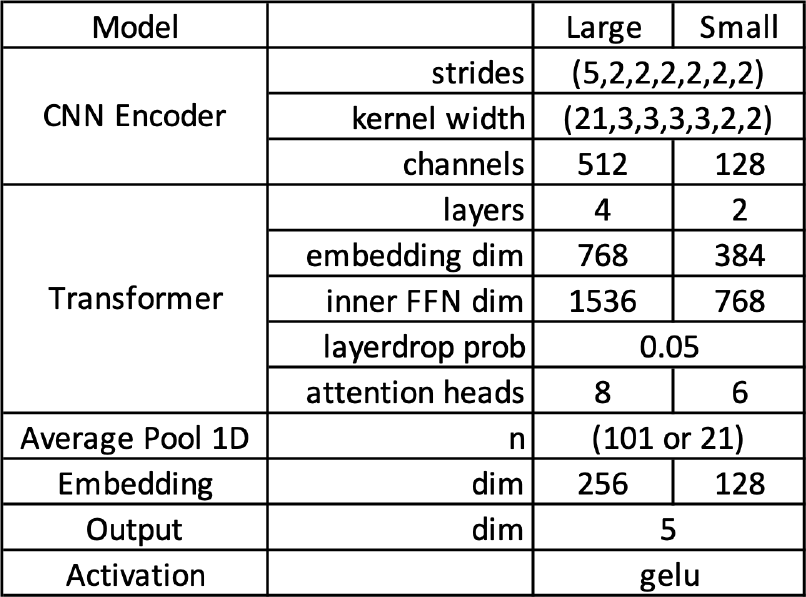
Transformer-based sleep staging model. Summary of parameters of model shown in Fig. 1.

We trained the larger variants of this model using input from both the EEG and EOG channels for a 101-epoch (or 50.5 minute) long input sequence (the ‘large’ model architecture described in Figure 2). The model was trained to minimize cross-entropy for 50 epochs (passes through the data) using an Adam optimizer. The learning rate ramped up linearly from 0.00001 to a peak of 0.000375 after the first ten passes. We retained the weights for the model corresponding to the training pass that achieved the lowest validation loss during the training run. After training we evaluated our models on the external validation data and the Sleep-EDF test data.

To understand the influence of different parts of the model architecture and training regime, we conducted a series of ablation studies using the external validation data to compare different parameters. All models used the same training hyperparameters described above except where noted. We tested a variety of configurations for the input data (EEG channel alone vs. both EEG+EOG together as input; 21-epoch vs. 101-epoch long input sequences; single nights of data per subject vs. multiple nights [where available]), and model architecture size (large vs. small models, see Figure 2 for details), for a total of 16 models varying in size from 3.9 million trainable parameters (23.4 MB on disk) to 28.4 million (129.2 MB on disk) trainable parameters. Models that took a 21-epoch input sequence were trained with a batch size of 64 and models taking the 101-epoch input sequence were trained with a batch size of 32.

## III. Results

Overall performance results from all sleep staging transformer model variants are displayed in Figure 4; more detailed confusion matrices for the large model are shown in Figure 3. This variant achieved exceptionally high performance on the internal validation data (87.7%) and continued to perform well on the external validation data (83.3% and 85% on Dreem-H and Dreem-O, respectively). This variant also performed well on the Sleep-EDF corpus (80.5%), which uses a slightly different recording montage than the training data (Cz-Fpz instead of C3 or C4). These results compare favorably with the reliability of human annotators (approx. 80-85%), as well as with other state-of-the-art systems (83.2% to 88.8% on Dreem data reported by Hanna & Flöel, 2023 [30]).

**Figure 3.**
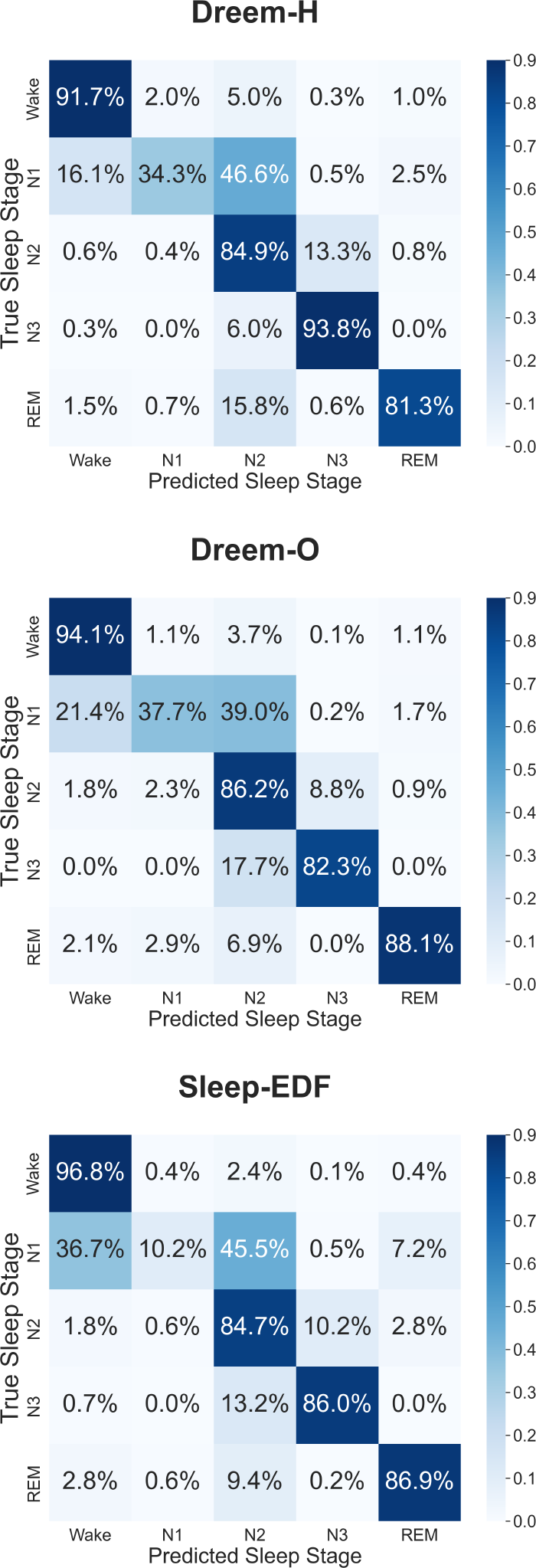
Performance Summaries by Sleep Stage. Confusion matrices show external validation performance from the top-performing ‘large’ model on three benchmark data sets. Predicted class is shown on ‘x-axis’. True sleep stages are shown on the ‘y-axis’. Since Sleep-EDF uses a different EEG referencing montage than the data used for training, this provides evidence of robust generalization on more out-of-domain data.

**Figure 4.**
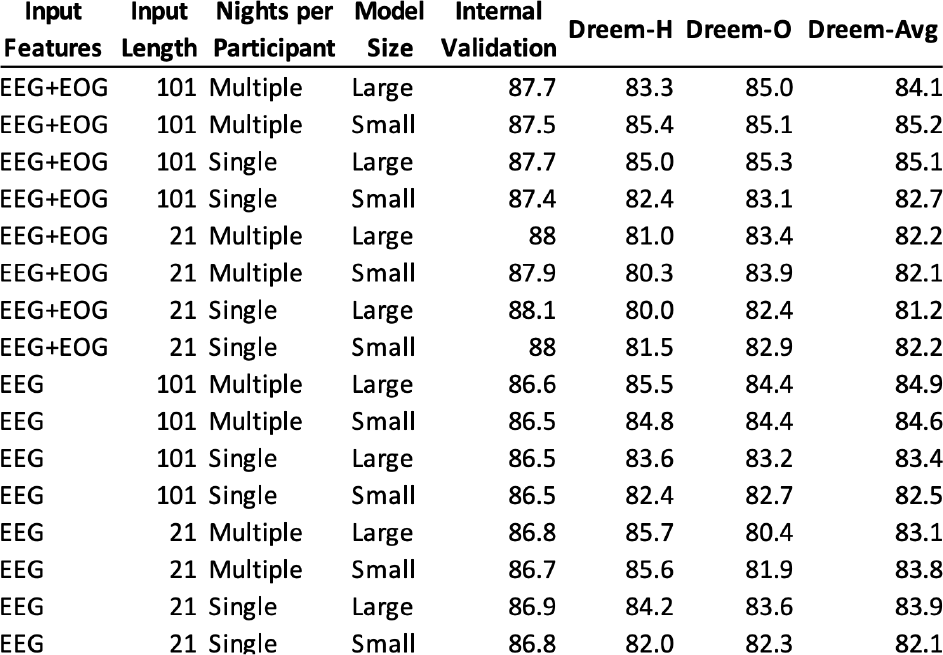
Overall performance of all model variants. Accuracies are shown as percentages.

Most model and input configurations achieved high internal validation performance and were able to match human expert performance (approx. 80-85% accuracy) on the Dreem validation data sets. Our best performing variant was the smaller architecture using EEG+EOG channels, multiple nights of sleep from participants where available, and longer 101-epoch input sequences. It is notable that this model slightly under-performed on our internal validation data relative to other models, but achieved better external validation performance. It is also notable that a minimal model (trained with the smallest input sequence length and model architecture) still performs well on the Dreem benchmark datasets (83.8%).

## IV. Discussion

Overall we found that these transformer models, which were originally developed solely for acoustic speech processing and language modeling, all proved adept at sleep-stage classification as well. They performed well testing on unseen data with a known character (i.e., internal validation, using data drawn from the same study as the training data^¶^) and also on unseen data with an unknown character (i.e., external validation, using data from a different study). Our results are nearly state-of-the-art using the Dreem and Sleep-EDF data sets for assessment (see comparisons in Hanna & Flöel, 2023

[30] and Yao & Liu, 2023 [31] for the Dreem and Sleep-EDF datasets, respectively). However, performance did drop on external validation data, and dropped even more when the recording montage differed (as for the Sleep-EDF data, which used a Cz-Fpz EEG derivation instead of the preferred C3-A2 or C4-A1). This generalization issue is an important problem to solve because it means that, currently, models will always need to be fine-tuned on the particular system or device on which they are deployed, which in turn implies a nontrivial amount of data collection beforehand. Ideally, one pretrained model would be able to perform equally well on traditional 10-20 locations and on nonstandard locations such as those used by commercial or at-home sleep monitoring wearables, without any fine-tuning to go from one to the other. In the companion paper appearing in this issue (Ogg & Coon, 2024) we report preliminary evidence that fine-tuning a foundation model could be one solution to this problem, since it vastly reduced the amount of data required for performant sleep stage classification.

It is curious that the large model variant performed better on the Dreem-O data set than the Dreem-H data set. The former is composed of slightly older individuals diagnosed with sleep apnea, while the latter is made up of healthy younger sleepers. We speculate that this difference in age and apnea state may have aligned better with the age distribution of our training data, which trended older and often involved studies targeting sleep apnea, possibly explaining the observed performance difference.

Smaller models generally performed as well as (or in some cases better than) comparable large models. This has important implications for deploying transformer-based sleep scoring models on mobile devices, for which storage space and processing power come at a premium (see Yao & Liu, 2023 [31] for some elegant work in this area). Encouragingly, the same high performance also held true for shorter input sequences, which could reduce the amount of processing power required. Discouragingly, the memory storage burden was high even for the small models (containing millions of high-precision weights), making them challenging to adapt for mobile use. Future work will explore shrinking and compressing these models (e.g., by using quantization and reduced precision to reduce their size).

We were motivated to study transformer model optimizations for sleep stage classification because of their suitability as foundation models for data-driven discovery down the line. Future work will examine self-supervised and masked training regimes for these tasks, which sidestep the bottleneck of needing human annotations and may be able to assimilate latent structure not captured by other sleep analysis techniques (including 5-stage scoring). Early results from one such study using this network as a foundation model can be found in a companion paper, Ogg & Coon 2024, which also appears in this issue of EMBC Proceedings. These foundation model variants point to the possibility of developing new, data-driven and objective sleep stage taxonomies that could more completely capture clinically informative facets of sleep. This could help address the recognized limitations of AASM sleep scoring guidelines [32], such as a “one-size-fits-all” lack of personalization (i.e., adjusting to the characteristics of individuals’ unique sleep electrophysiology), age-wise scoring biases, disease state interactions, and other recognized phenomena such as the cyclic alternating pattern (CAP) [33] that are not captured by current scoring criteria. In pursuing these new methods it will be critical to anchor any study of their utility to the baselines established here, to tie them back to established methods for comparison. This work hence lays a foundation that may unlock myriad other health applications in the future.

## Acknowledgment

This study was supported by the Johns Hopkins University Applied Physics Laboratory under an internal research & development grant. The Cleveland Family Study (CFS) was supported by grants from the National Institutes of Health (HL46380, M01 RR00080-39, T32-HL07567, RO1-46380).

The National Heart, Lung, and Blood Institute provided funding for the ancillary MrOS Sleep Study, “Outcomes of Sleep Disorders in Older Men,” under the following grant numbers: R01 HL071194, R01 HL070848, R01 HL070847, R01 HL070842, R01 HL070841, R01 HL070837, R01 HL070838, and R01 HL070839. The Multi-Ethnic Study of Atherosclerosis (MESA) Sleep Ancillary study was funded by NIH-NHLBI Association of Sleep Disorders with Cardiovascular Health Across Ethnic Groups (RO1 HL098433). MESA is supported by NHLBI funded contracts HHSN268201500003I, N01-HC-95159, N01-HC-95160, N01-HC-95161, N01-HC-95162, N01-HC-95163, N01-HC-95164, N01-HC-95165, N01-HC-95166, N01-HC-95167, N01-HC-95168 and N01-HC-95169 from the National Heart, Lung, and Blood Institute, and by cooperative agreements UL1-TR-000040, UL1-TR-001079, and UL1-TR-001420 funded by NCATS. NCH Sleep DataBank was supported by the National Institute of Biomedical Imaging and Bioengineering of the National Institutes of Health under Award Number R01EB025018. The Sleep Heart Health Study (SHHS) was supported by National Heart, Lung, and Blood Institute cooperative agreements U01HL53916 (University of California, Davis), U01HL53931 (New York University), U01HL53934 (University of Minnesota), U01HL53937 and U01HL64360 (Johns Hopkins University), U01HL53938 (University of Arizona), U01HL53940 (University of Washington), U01HL53941 (Boston University), and U01HL63463 (Case Western Reserve University). The Study of Osteoporotic Fractures (SOF) was supported by National Insti-tutes of Health grants (AG021918, AG026720, AG05394, AG05407, AG08415, AR35582, AR35583, AR35584, RO1 AG005407, R01 AG027576-22, 2 R01 AG005394-22A1, 2 RO1 AG027574-22A1, HL40489, T32 AG000212-14). This Wisconsin Sleep Cohort Study was supported by the U.S. National Institutes of Health, National Heart, Lung, and Blood Institute (R01HL62252), National Institute on Aging (R01AG036838, R01AG058680), and the National Center for Research Resources (1UL1RR025011). The National Sleep Research Resource was supported by the U.S. National Institutes of Health, National Heart Lung and Blood Institute (R24 HL114473, 75N92019R002)

## Footnotes

† traditionally sleep is scored using polysomnography (PSG), an aggregate of EEG, electro-oculography (EOG), and electromyography (EMG), but most information is in the EEG’s brain signals and many automated techniques perform well using EEG alone [1]

‡ European Data Format. EDF is an industry standard file format for polysomnographic records.

§ values were selected to vary the amount of sleep architecture history the model could assess at a time; the long sequence model captures most of a sleep cycle, while the short variant presumably cannot benefit from this informative background context.

¶ both the training data used for training the model, and the held out test data used to evaluate performance during model training, were drawn from the same source data corpus (for example, the MrOS study), but did NOT overlap.

## Notes

### Competing Interest Statement

The authors have declared no competing interest.

### Summary of Updates

We caught an error in the reported learning rate schedules, and have updated accordingly. Narrative refinements and additional citations have been added.

## References

[1] Huy Phan and Kaare Mikkelsen. Automatic sleep staging of eeg signals: recent development, challenges, and future directions. Physiological Measurement, 43(4):04TR01, 2022. doi:10.1088/1361-6579/ac6049.

[2] Kales A Rechtschaffen A (eds). A Manual of Standardized Terminology, Techniques and Scoring System for Sleep Stages of Human Subjects. BIS/BRI University of California, Los Angeles, 1968.

[3] Michael H Silber, Sonia Ancoli-Israel, Michael H Bonnet, Sudhansu Chokroverty, Madeleine M Grigg-Damberger, Max Hirshkowitz, Sheldon Kapen, Sharon A Keenan, Meir H Kryger, Thomas Penzel, et al. The visual scoring of sleep in adults. Journal of Clinical Sleep Medicine, 3(02):121–131, 2007. doi:10.5664/jcsm.26814.

[4] Henri Korkalainen, Juhani Aakko, Sami Nikkonen, Samu Kainulainen, Akseli Leino, Brett Duce, Isaac O Afara, Sami Myllymaa, Juha Töyräs, and Timo Leppänen. Accurate deep learning-based sleep staging in a clinical population with suspected obstructive sleep apnea. IEEE journal of biomedical and health informatics, 24(7):2073–2081, 2019.

[5] William G Coon and Naresh M Punjabi. Automatic sleep staging using a small-footprint sensor array and recurrent-convolutional neural networks. In 2021 10th International IEEE/EMBS Conference on Neural Engineering (NER), pages 1144–1147. IEEE, 2021. doi:10.1109/NER49283.2021.9441432.

[6] Akara Supratak, Hao Dong, Chao Wu, and Yike Guo. Deepsleepnet: A model for automatic sleep stage scoring based on raw single-channel eeg. IEEE Transactions on Neural Systems and Rehabilitation Engineering, 25(11):1998–2008, 2017.

[7] Mathias Perslev, Sune Darkner, Lykke Kempfner, Miki Nikolic, Poul Jørgen Jennum, and Christian Igel. U-sleep: resilient highfrequency sleep staging. NPJ Digital Medicine, 4(1):72, 2021. doi:10.1038/s41746-021-00440-5.

[8] Ziyu Jia, Youfang Lin, Jing Wang, Ronghao Zhou, Xiaojun Ning, Yuanlai He, and Yaoshuai Zhao. Graphsleepnet: Adaptive spatial-temporal graph convolutional networks for sleep stage classification. In IJCAI, volume 2021, pages 1324–1330, 2020.

[9] Alexander Neergaard Olesen, Poul Jørgen Jennum, Emmanuel Mignot, and Helge Bjarup Dissing Sorensen. Automatic sleep stage classification with deep residual networks in a mixed-cohort setting. Sleep, 44(1):zsaa161, 2021.

[10] Yinghao Li, Zhenghui Gu, Zichao Lin, Zhuliang Yu, and Yuanqing Li. An automatic sleep staging model combining feature learning and sequence learning. In 2020 12th International Conference on Advanced Computational Intelligence (ICACI), pages 419–425. IEEE, 2020.

[11] Emadeldeen Eldele, Zhenghua Chen, Chengyu Liu, Min Wu, Chee-Keong Kwoh, Xiaoli Li, and Cuntai Guan. An attention-based deep learning approach for sleep stage classification with single-channel eeg. IEEE Transactions on Neural Systems and Rehabilitation Engineering, 29:809–818, 2021.

[12] Antoine Guillot and Valentin Thorey. Robustsleepnet: Transfer learning for automated sleep staging at scale. IEEE Transactions on Neural Systems and Rehabilitation Engineering, 29:1441–1451, 2021.

[13] Huy Phan, Kaare Mikkelsen, Oliver Y Chén, Philipp Koch, Alfred Mertins, and Maarten De Vos. Sleeptransformer: Automatic sleep staging with interpretability and uncertainty quantification. IEEE Transactions on Biomedical Engineering, 69(8):2456–2467, 2022.

[14] Yang Dai, Xiuli Li, Shanshan Liang, Lukang Wang, Qingtian Duan, Hui Yang, Chunqing Zhang, Xiaowei Chen, Longhui Li, Xingyi Li, et al. Multichannelsleepnet: A transformer-based model for automatic sleep stage classification with psg. IEEE Journal of Biomedical and Health Informatics, 2023.

[15] Jathurshan Pradeepkumar, Mithunjha Anandakumar, Vinith Kugathasan, Dhinesh Suntharalingham, Simon L Kappel, Anjula C De Silva, and Chamira US Edussooriya. Towards interpretable sleep stage classification using cross-modal transformers. arXiv preprint arXiv:2208.06991, 2022.

[16] Nicolas Decat, Jasmine Walter, Zhao H Koh, Piengkwan Sribanditmongkol, Ben D Fulcher, Jennifer M Windt, Thomas Andrillon, and Naotsugu Tsuchiya. Beyond traditional sleep scoring: Massive feature extraction and data-driven clustering of sleep time series. Sleep Medicine, 98:39–52, 2022. doi:10.1016/j.sleep.2022.06.013.

[17] Wei-Ning Hsu, Benjamin Bolte, Yao-Hung Hubert Tsai, Kushal Lakhotia, Ruslan Salakhutdinov, and Abdelrahman Mohamed. Hubert: Self-supervised speech representation learning by masked prediction of hidden units. IEEE/ACM Transactions on Audio, Speech, and Language Processing, 29:3451–3460, 2021. doi:10.1109/TASLP.2021.3122291.

[18] Jacob Devlin, Ming-Wei Chang, Kenton Lee, and Kristina Toutanova. Bert: Pre-training of deep bidirectional transformers for language understanding. arXiv preprint arXiv:1810.04805, 2018. doi:10.48550/arXiv.1810.04805.

[19] Guo-Qiang Zhang, Licong Cui, Remo Mueller, Shiqiang Tao, Matthew Kim, Michael Rueschman, Sara Mariani, Daniel Mobley, and Susan Redline. The national sleep research resource: towards a sleep data commons. Journal of the American Medical Informatics Association, 25(10):1351–1358, 2018. doi:10.1093/jamia/ocy064.

[20] Xiaoli Chen, Rui Wang, Phyllis Zee, Pamela L Lutsey, Sogol Javaheri, Carmela Alcántara, Chandra L Jackson, Michelle A Williams, and Susan Redline. Racial/ethnic differences in sleep disturbances: the multi-ethnic study of atherosclerosis (mesa). Sleep, 38(6):877–888, 2015. doi:10.5665/sleep.4732.

[21] Terri Blackwell, Kristine Yaffe, Sonia Ancoli-Israel, Susan Redline, Kristine E Ensrud, Marcia L Stefanick, Alison Laffan, Katie L Stone, and Osteoporotic Fractures in Men (MrOS) Study Group. Association of sleep characteristics and cognition in older community-dwelling men: the mros sleep study. Sleep, 34(10):1347–1356, 2011. doi:10.5665/SLEEP.1276.

[22] Stuart F Quan, Barbara V Howard, Conrad Iber, James P Kiley, F Javier Nieto, George T O’Connor, David M Rapoport, Susan Redline, John Robbins, Jonathan M Samet, et al. The sleep heart health study: design, rationale, and methods. Sleep, 20(12):1077–1085, 1997. doi:10.1093/sleep/20.12.1077.

[23] Terry Young, Mari Palta, Jerome Dempsey, Paul E Peppard, F Javier Nieto, and K Mae Hla. Burden of sleep apnea: rationale, design, and major findings of the wisconsin sleep cohort study. WMJ: official publication of the State Medical Society of Wisconsin, 108(5):246, 2009. doi:PMID: 19743755 PMCID: PMC2858234.

[24] Harlin Lee, Boyue Li, Shelly DeForte, Mark L Splaingard, Yungui Huang, Yuejie Chi, and Simon L Linwood. A large collection of real-world pediatric sleep studies. Scientific Data, 9(1):421, 2022. doi:10.1038/s41597-022-01545-6.

[25] Adam P Spira, Terri Blackwell, Katie L Stone, Susan Redline, Jane A Cauley, Sonia Ancoli-Israel, and Kristine Yaffe. Sleep-disordered breathing and cognition in older women. Journal of the American Geriatrics Society, 56(1):45–50, 2008. doi:10.1111/j.1532-5415.2007.01506.x.

[26] Susan Redline, Peter V Tishler, Tor D Tosteson, John Williamson, Kenneth Kump, Ilene Browner, Veronica Ferrette, and Patrick Krejci. The familial aggregation of obstructive sleep apnea. American journal of respiratory and critical care medicine, 151 (3):682–687, 1995. doi:10.1164/ajrccm.151.3.7881656.

[27] Antoine Guillot, Fabien Sauvet, Emmanuel H During, and Valentin Thorey. Dreem open datasets: Multi-scored sleep datasets to compare human and automated sleep staging. IEEE transactions on neural systems and rehabilitation engineering, 28(9):1955–1965, 2020. doi:10.1109/TNSRE.2020.3011181.

[28] Bob Kemp, Aeilko Zwinderman, B Tuk, H Kamphuisen, and J Oberyé. Sleep-edf database expanded. Physionet org, 2018. doi:10.13026/C2X676.

[29] Ashish Vaswani, Noam Shazeer, Niki Parmar, Jakob Uszkoreit, Llion Jones, Aidan N Gomez, Łukasz Kaiser, and Illia Polosukhin. Attention is all you need. Advances in neural information processing systems, 30, 2017.

[30] Jevri Hanna and Agnes Flöel. An accessible and versatile deep learning-based sleep stage classifier. Frontiers in Neuroinformatics, 17:1086634, 2023. doi:10.3389/fninf.2023.1086634.

[31] Zongyan Yao and Xilin Liu. A CNN-transformer deep learning model for real-time sleep stage classification in an energy-constrained wireless device. In 2023 11th International IEEE/EMBS Conference on Neural Engineering (NER), pages 1–4. IEEE, 2023. doi:ieeexplore.ieee.org/abstract/document/10123825/.

[32] Sari-Leena Himanen and Joel Hasan. Limitations of rechtschaffen and kales. Sleep medicine reviews, 4(2):149–167, 2000. doi:10.1053/smrv.1999.0086.

[33] Mario Giovanni Terzano and Liborio Parrino. Origin and significance of the cyclic alternating pattern (cap). Sleep medicine reviews, 4(1):101–123, 2000. doi:10.1053/smrv.1999.0083.

